# Endothelial features along the pulmonary vascular tree in chronic thromboembolic pulmonary hypertension: distinctive or shared facets?

**DOI:** 10.1101/2024.02.27.582290

**Authors:** Janne Verhaegen, Lynn Willems, Allard Wagenaar, Ruben Spreuwers, Nessrine Dahdah, Lucia Aversa, Tom Verbelen, Marion Delcroix, Rozenn Quarck

## Abstract

Chronic thromboembolic pulmonary hypertension (CTEPH) is a rare complication of pulmonary embolism, characterized by the presence of organized fibro-thrombotic material that partially or fully obstructs the lumen of large pulmonary arteries, microvasculopathy, and enlargement of the bronchial systemic vessels. The precise mechanisms underlying CTEPH remain unclear. However, defective angiogenesis and altered pulmonary arterial endothelial cell (PAEC) function may contribute to disease progression. Despite the observation of differences in histological features, shear stress and ischemia along the pulmonary vascular tree, the potential contribution of PAEC phenotype and function to these disparate aspects remains unexplored. Based on these observations, we postulated that angiogenic capacities and endothelial barrier function may contribute to disparities in histological features observed along the pulmonary vascular tree.

We thus explored the histological characteristics of the pulmonary vascular tree using pulmonary arterial lesions obtained during pulmonary endarterectomy (PEA). We focused on the angiogenic vascular endothelial growth factor (VEGF)-A/VEGF receptor-2 (VEGFR2) axis and collagen 15A1 (COL15A1), a potential marker of endothelial cells of the systemic circulation. Concurrently, we examined *in vitro* angiogenic properties and barrier function of PAECs derived from large and (sub)-segmental pulmonary arterial lesions. (Sub)-segmental pulmonary arterial lesions were abundantly recanalized by neovessels, paralleled by an enriched expression of VEGFR2. VEGF-A expression was more pronounced in large pulmonary arterial lesions. Nevertheless, no significant difference was discerned in *in vitro* angiogenic capacities and barrier integrity of PAECs isolated from large and (sub)- segmental pulmonary arterial lesions. Importantly, our findings revealed the presence of endothelial cells (CD31^+^) expressing COL15A1, as well as CD31^+^ cells that did not express COL15A1. This suggests that endothelial cells from both systemic and pulmonary circulation contribute to lesion recanalization.

Despite disparate *in situ* angiogenic cues in VEGF-A/VEGFR2 axis between large and (sub)- segmental pulmonary arterial lesions in CTEPH, *in vitro* angiogenic capacities and barrier function remain unaltered.

## INTRODUCTION

Chronic thromboembolic pulmonary hypertension (CTEPH) is a rare complication of massive or recurrent pulmonary embolism (1–4). The condition is characterized by the presence of intra-luminal unresolved thrombi associated with fibrous stenosis that partially or completely obstructs the large pulmonary arteries. This eventually results in increased pulmonary vascular resistance (PVR), progressive pulmonary hypertension (PH), and right ventricular (RV) failure, if left untreated (2,5,6). Additionally, CTEPH is associated with progressive microvasculopathy, encompassing remodeling and obstruction of pulmonary microvessels in perfused and non-perfused areas (5,7,8), and presence of bronchial systemic-to-pulmonary anastomoses (9). The treatment of choice for operable patients remains surgical removal of the fibro-thrombotic material, by pulmonary endarterectomy (PEA) (10).

Various risk factors increasing the likelihood of CTEPH development, including venous thromboembolism, inflammatory diseases, splenectomy, non-O blood group, cancer, insufficient anticoagulation, increased antiphospholipid antibodies and lupus anticoagulant (5,11), have been identified. However, the precise sequence of cellular and molecular mechanisms leading to sustained fibro-thrombotic obstruction remains to be fully understood. Several hypotheses have been put forth the underlying mechanisms of CTEPH. These include i) dysregulated fibrinolysis (12–15); ii) inflammatory thrombosis (12,16–22); iii) activation of pulmonary artery endothelial (PAECs), smooth muscle (PASMCs), and progenitor cells (23–26); and iv) deficient angiogenesis (27–32). Prior research has indicated that angiogenesis plays a pivotal role in thrombus resolution and neovascularization, which is associated with the outcome of CTEPH (19,33). PAECs isolated from PEA material were observed to exhibit reduced angiogenic properties compared to lung donor PAECs, suggesting their potential contribution to CTEPH pathogenesis (34). Moreover deficient angiogenesis was shown to play a crucial role in the progression to CTEPH in a unique rabbit model (35). The two-faced nature of angiogenesis in CTEPH is discussed in great detail in a recent review (36). The integrity of the pulmonary endothelium as a barrier is of paramount importance in preventing the infiltration of inflammatory cells, deemed to contribute to the progression of CTEPH.

It is noteworthy that previous histological analyses of pulmonary vascular lesions have revealed discrepancies in cellular and extracellular matrix composition along the pulmonary vascular tree, with considerable variations observed in the extend of neovascularization across the lesion (16,17,19). Accordingly, proximal lesions are predominantly parietalized material, whereas distal lesions material appear to be more occlusive. It has been proposed that these disparities may be attributed to variations in flow dynamics, *e.g.* high-shear stress upstream of the occlusion, in contrast to with low-shear stress and disrupted blood flow downstream of the occlusion (5). Additionally, angiogenic signaling markers, including vascular endothelial growth factor A (VEGF-A) and vascular endothelial growth factor receptor 2 (VEGFR2) exhibit are differential expression in fresh versus organized thrombi, derived from PEA material (37). Interestingly, Schupp *et al.* identified COL15A1 as a potential marker of a subpopulation of systemic endothelial cells, not expressed by pulmonary endothelial cells (38). However, it remains unclear whether the *in situ* expression patterns of endothelial and/or angiogenic markers along the pulmonary vascular tree can be attributed to angiogenic behavior or barrier function of PAECs.

Thus, we hypothesized that *in vitro* PAEC phenotype and function may vary according to their position along the pulmonary vascular lesion tree in CTEPH patients. The present study aimed to explore histological characteristics of pulmonary arterial lesions collected during PEA procedures and to investigate *in vitro* angiogenic capacities and barrier function of PAECs according to their location along the pulmonary vascular tree in CTEPH patients.

## MATERIALS AND METHODS

### Study population

This study includes pulmonary vascular material from eight patients with CTEPH, who underwent PEA at University Hospital Leuven between August 2018 and April 2023. The study protocol was approved by the Institutional Ethics Committee of the University Hospital Leuven (S57114) and participants gave written informed consent preoperatively.

### Tissue collection

Obstructive material was collected at the time of PEA and the pulmonary vascular tree was reconstructed with the help of the surgeon. A specimen is shown in **Figure 1**. A piece from the large (upstream pulmonary vascular occlusion) and (sub)-segmental (downstream pulmonary vascular occlusion) pulmonary arteries, free of thrombotic material, was collected separately in endothelial growth medium MV2 (PromoCell) supplemented with 100 U.mL^-1^ penicillin, 100 µg.ml^-1^ 100 µg.mL^-1^ streptomycin and 1.25 µg.mL^-1^ amphotericin B (Life Technologies).

**Figure 1:**
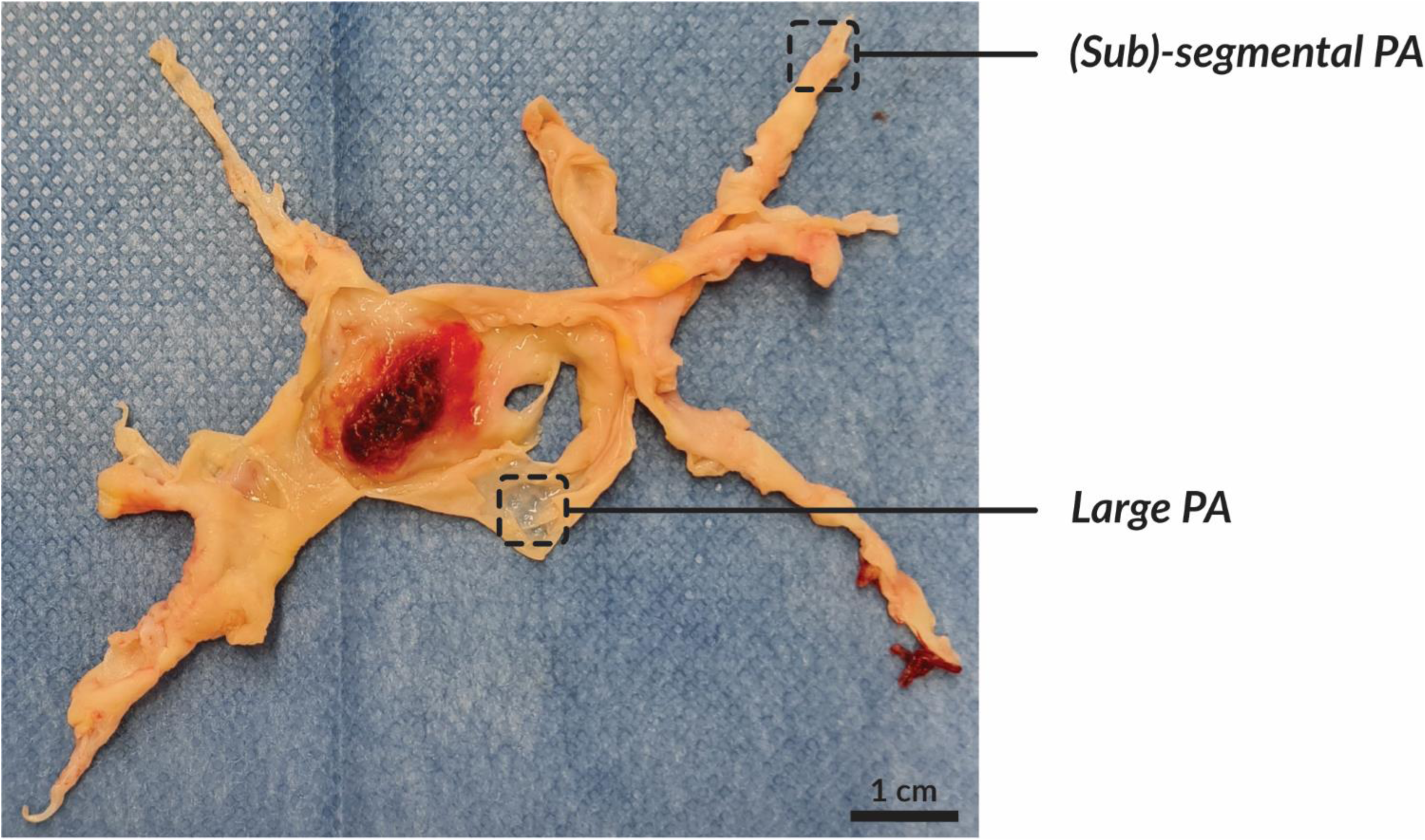
Representative resection specimen obtained during PEA. Fibro-thrombotic material obstructing pulmonary arteries is divided into large pulmonary artery, upstream of the obstruction by thrombotic material, and (sub)-segmental pulmonary artery, downstream of the obstruction by thrombotic material. PA, pulmonary artery; PEA, pulmonary endarterectomy.

### Histological assessment

Large and (sub)-segmental pulmonary arterial lesions were fixed in 4% paraformaldehyde and embedded in paraffin blocks. Five-µm thick serial sections were deparaffinized and rehydrated. Collagen fibers, fibrin and muscle fibers were stained with Masson’s Trichrome. Briefly, slides were fixed in Bouin’s solution overnight and stained with Weigert’s iron hematoxylin, Biebrich scarlet-acid fuchsin, phosphotungstic/phosphomolybdic acid, aniline blue and acetic acid working solutions, respectively. To examine endothelial markers, the presence of smooth muscle cells and angiogenic signaling, immunolabeling was performed with mouse monoclonal antibodies against CD31 (1:50, clone JC70A; Dako), alpha smooth muscle actin (αSMA; 1:750; clone 1A4; Dako) and VEGF-A (1:100; clone VG-1; Abcam), rabbit monoclonal antibodies against VEGFR2 (1:200, clone 55b11, Cell Signalling) and rabbit polyclonal antibodies against COL15A1 (1:200, Invitrogen). Immunolabelling was further revealed using horseradish peroxidase-conjugated goat anti-mouse (1:1, Abcam) secondary antibodies for CD31, αSMA and VEGF-A, and horseradish peroxidase-conjugated goat anti-rabbit (1:1, Abcam) secondary antibodies for VEGFR2 and COL15A1, and DAB substrate kit (Abcam) according to the manufacturer’s instructions. Cell nuclei were counterstained with hematoxylin. Images were captured using Axio Scan.Z1 (Zeiss) at 20x magnification. DAB-positive regions (positive pixels per total number of pixels) were quantified using the open-source software QuPath (https://qupath.github.io). Only patients with at least one section of both large and (sub)- segmental pulmonary arterial lesions were included for quantification. For technical reasons, patient cohort was reduced to n=4 for CD31, COL15A1 and αSMA, and n=6 for VEGF-A and VEGFR2.

### Isolation and purification of PAECs

PAECs were isolated from large and (sub)-segmental pulmonary arterial lesions by collagenase I digestion (1 mg.mL^-1^, Gibco) for 20 and 30 min at 37°C, respectively (24,39). Cells were seeded onto gelatine-coated flasks (2 mg/mL; Sigma-Aldrich) in endothelial growth medium MV2 (PromoCell) supplemented with 100 U.mL^-1^ penicillin, 100 µg.mL^-1^ streptomycin and 1.25 µg.mL^-1^ amphotericin B (Life Technologies) and maintained in culture at 37°C, 5% CO2 and 95% humidity. To obtain homogeneous populations of PAECs, cells were purified by immunomagnetic separation using anti-CD31 monoclonal antibody-labeled beads, according to the manufacturer’s instructions (Miltenyi Biotec). Before each experiment, PAECs were starved with basal endothelial medium MV2 (Promocell) containing 3.2% endothelial growth medium, supplemented with 100 U.mL^-1^ penicillin, 100 µg.mL^-1^ streptomycin and 1.25 µg.mL^-1^ amphotericin B. All experiments were performed with PAECs between passages 3 and 8.

### Cell phenotyping

PAECs (3.000 cells/chamber) were seeded onto fibronectin-coated (10 µg.mL^-1^; R&D Systems) 4-chamber slides (Nunc). PAECs were immunolabeled with primary mouse monoclonal antibodies against CD31 (1:25; clone JC70A; Dako) and vWF (1:50; clone F8/86; Dako), rabbit monoclonal antibodies against VE-cadherin (1:600; clone D87F2; Cell Signaling Technology), and Alexa Fluor-594 goat anti-mouse (1:2000; Life Technologies) for CD31 and vWF and Alexa Fluor-488 goat anti-rabbit (1:2000; Life Technologies) for VE-cadherin secondary antibodies, as previously described (40). Cell nuclei were stained with DAPI. Endothelial scavenger receptor labeling was performed with dil-Ac-LDL (1:20; Tebubio), as previously described (39–41). All fluorescence images were captured at a 40x magnification with an Olympus BX-UCB microscope equipped with X-Cite series 120Q (Excelitas Technologies).

### Migration assay

PAECs were seeded onto gelatin-coated 2-well culture inserts (Ibidi, 81176) at a density of 7,500 cells/well (42). Confluent PAECs were starved for 6 h, inserts were removed, and PAECs were placed in fresh endothelial growth medium and imaged every 2 h for 20 h using EVOS Cell Imaging System (Thermo Fisher Scientific) equipped with an EVOS Onstage Incubator (37°C, 5% CO2 and 95% humidity; Thermo Fisher Scientific). At each time point, the gap area was measured using ImageJ and subtracted from the initial gap area (baseline), plotted over time, and the area under the curve (AUC) was calculated, which corresponds to the migration rate in mm^2^.h^-1^.

### Tube formation assay

Angiogenesis µ-slides (Ibidi, 81506) were filled with Matrigel^©^ (BD Bioscience) and incubated at 37°C for 30 min, as described elsewhere (42). PAECs were starved overnight, trypsinized, resuspended in endothelial growth medium and seeded on Matrigel^©^ at a density of 5,000 cells/well. After a 4-h incubation at 37°C, cells were imaged using the EVOS Cell Imaging System. The number of nodes, junctions, branches and tubes was quantified in triplicate using the ‘Angiogenesis Analyzer’ plug-in for ImageJ, developed by Gilles Carpentier (43).

### Barrier function assay

PAECs (10,000 cells/insert) suspended in growth medium were seeded onto gelatin-coated inserts of 24-well Transwells (Corning)(42,44). After 24 h, the cells were starved overnight, and starvation medium containing 0.1 mg.mL^-1^ fluorescein isothiocyanate (FITC)-labeled bovine serum albumin (BSA; Sigma Aldrich) was added to the upper chamber, while the lower chamber was filled with starvation medium. Leakage of FITC-BSA across PAECs monolayer was assessed by collecting medium from the lower chamber at 30 min, 1, 2, and 4 h. Absorbance was measured (excitation 488 nm; emission 520 nm) using Synergy H1 hybrid reader (Biotek) and Gen5 Software (Biotek). The leakage of FITC-BSA across the PAECs monolayer was plotted against time and AUC was calculated and corresponds to the leakage rate in µg.mL^-1^.h^-1.^

### Statistical analysis

Statistical analyses were performed with Graphpad Prism (version 9; GraphPad Software Inc.). Normality was tested using Shapiro-Wilk test. Differences between two groups were analyzed using paired Student or Wilcoxon t-test, and 2-way multiple comparison ANOVA followed by post-hoc Šídák’s test. A p-value of <0.05 was considered statistically significant. Data are expressed as median (25^th^-75^th^ interquartile range) in graphs and as mean ± SD or otherwise indicated in the text.

## RESULTS

### Histomorphometry of large and (sub)-segmental pulmonary arterial lesions from CTEPH patients

Histological features along the pulmonary vascular tree, collagen and muscle fiber content and expression of endothelial (CD31, COL15A1), smooth muscle (αSMA) and angiogenic markers (VEGF-A and VEGFR2) were examined in sections of large and (sub)-segmental pulmonary arterial lesions (**Figure 2, 3**). Large pulmonary arterial lesions consist of media, with several layers of smooth muscle cells, and neointima with spindle-shaped cells, possibly myofibroblast-like cells (**Figure 2**). (sub)-segmental pulmonary arterial lesions (**Figure 3**) consist of a large amount of dense collagen fibers, with numerous recanalizing vessels surrounded by multiple layers of smooth muscle cells. CD31-positive recanalizing vessels are more abundant in (sub)-segmental (**Figure 3**) compared to large pulmonary arteries (**Figure 2**), as evidenced by a significantly larger CD31-positive area within (sub)-segmental compared to large pulmonary arterial lesions (1.65 ± 1.17% vs. 5.60 ± 1.11%; p = 0.0163; **Figure 4**). Interestingly, some CD31-positive recanalizing vessels expressed COL15A1, a marker of ECs from the systemic circulation, whereas a substantial proportion did not (**Figure 3; Figure S1**). This suggests that vessels recanalizing fibrothrombotic lesions may originate from either the bronchial and/or pulmonary circulation. Notably, CD31-positive cells lining the lumen of obstructed (sub)-segmental pulmonary arterial lesions did not express COL15A1 (**Figure S2**). Surprisingly, within the large pulmonary arterial wall, CD31-negative myofibroblast-like cells do express COL15A1 (**Figure 2**). Overall, the COL15A1-positive area tended to be higher in large compared to (sub)-segmental pulmonary arterial lesions (18.33 ± 10.70% vs. 4.72 ± 3.52%; p = 0.0525; **Figure 4**), probably because of the abundant presence of COL15A1- positive myofibroblast-like cells within large pulmonary arterial lesions. Furthermore, myofibroblast-like cells showed αSMA expression within the large pulmonary artery lesions (**Figure 2**). In contrast, αSMA expression was distributed throughout the lesions obstructing (sub)-segmental pulmonary arteries and in the tissue surrounding recanalising vessels (**Figure 3**). No significant difference in αSMA expression was observed between large and (sub)- segmental pulmonary arterial lesions (19.56 ± 12.79% vs. 26.68 ± 13.35; p = 0.4799; **Figure 4**). VEGF-A was expressed in both large and (sub)-segmental pulmonary arterial lesions, with significantly higher expression in large pulmonary arterial lesions (25.96 ± 16.61% vs. 7.72 ± 7.93%; p = 0.0260; **Figure 4**). In addition, neointimal myofibroblast-like cells and CD31- negative cells lining the lumen of large pulmonary arterial lesions expressed VEGF-A (**Figure 2**), whereas within (sub)-segmental pulmonary arterial lesions, CD31-positive cells lining recanalizing vessels did not express VEGF-A (**Figure 3**). Finally, VEGFR2 was expressed by myofibroblast-like cells in large pulmonary arterial lesions (**Figure 2**); in (sub)-segmental pulmonary arterial lesions VEGFR2 is expressed by αSMA-positive cells of recanalizing vessels (**Figure 3**). However, no significant difference in VEGFR2 expression was observed between large and (sub)-segmental pulmonary arterial lesions (13.51% [7.29-18.56%] vs. 17.74% [16.94-30.97%], median [interquartile range]; p = 0.2188; **Figure 4**).

**Figure 2:**
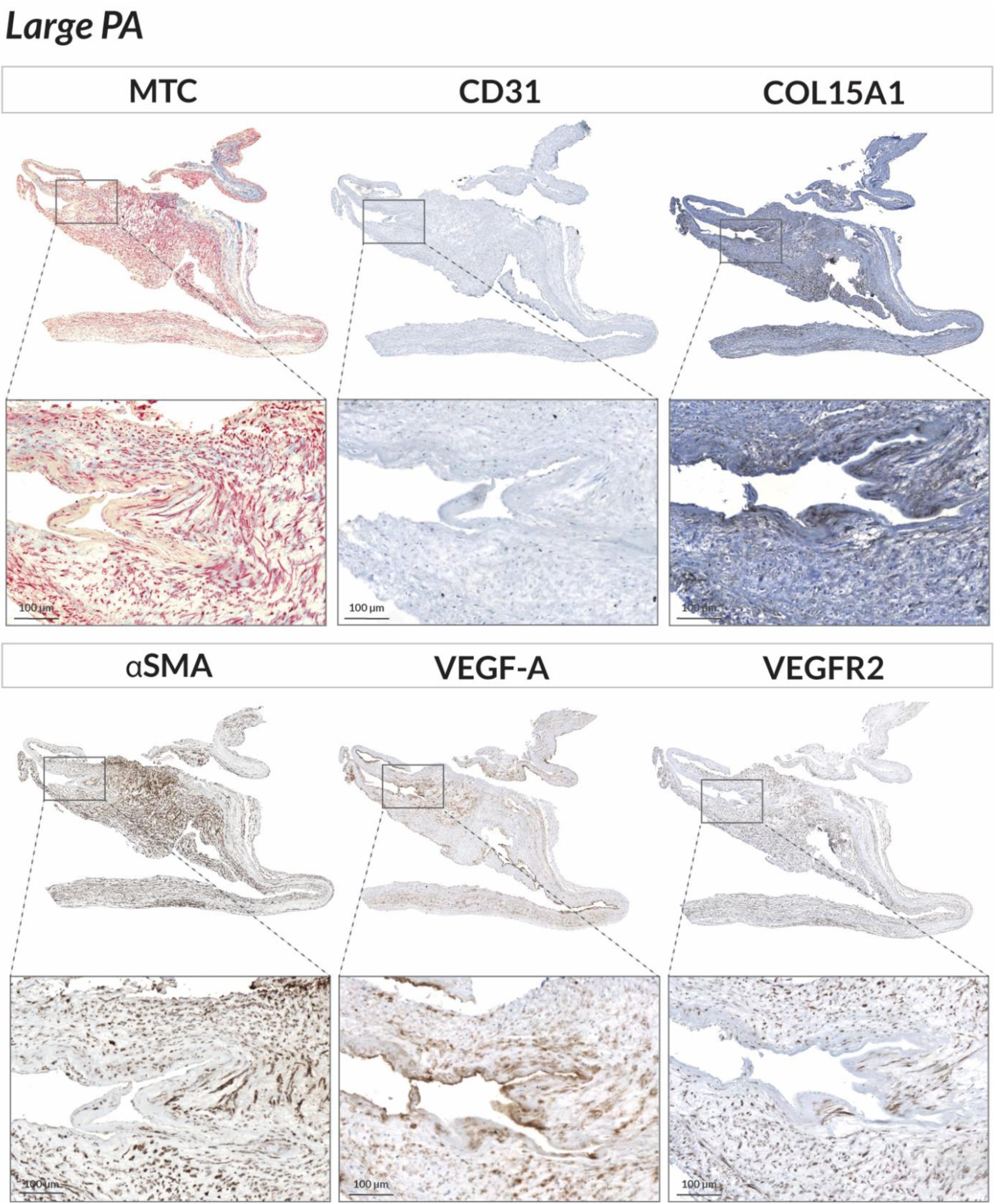
Histological evaluation of large pulmonary arterial lesions of CTEPH patients. Large pulmonary artery tissues were stained with Masson’s trichrome stain (collagen, blue; muscle fibers, red) and immunolabeled using CD31, COL15A1, αSMA, VEGF-A, and VEGFR2 antibodies. In total, PEA material of eight patients was evaluated and similar lesions were observed. Scale bar = 100 µm. PA, pulmonary artery; PEA, pulmonary endarterectomy; MTC, Masson trichrome; αSMA, alpha smooth muscle actin; VEGF-A, vascular endothelial growth factor A; VEGFR2, vascular endothelial growth factor receptor 2.

**Figure 3:**
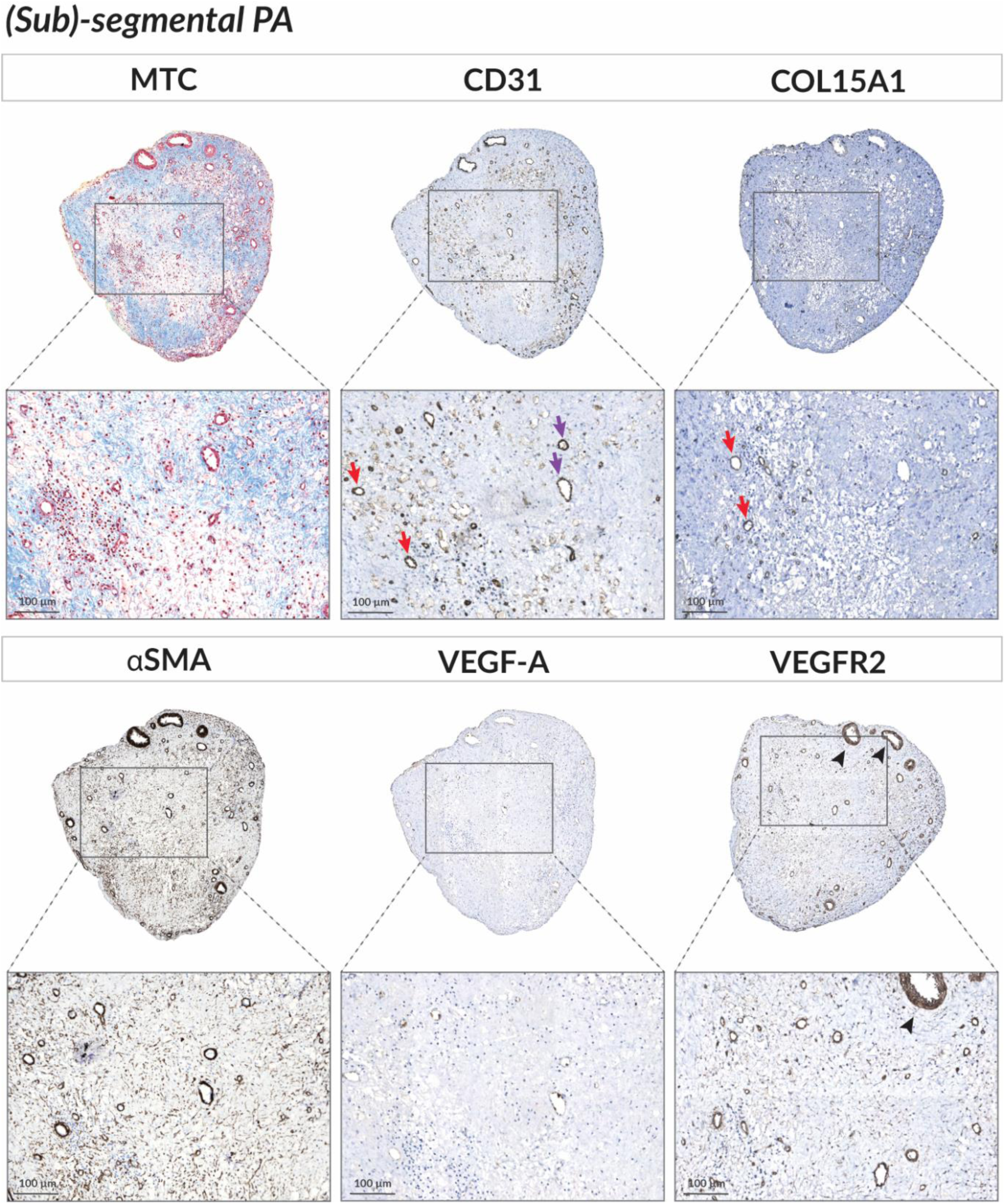
Histological evaluation of (sub)-segmental pulmonary arterial lesions of CTEPH patients. (Sub)-segmental pulmonary arterial lesions were stained with Masson’s trichrome stain (collagen, blue; muscle fibers, red) and immunolabeled using CD31, COL15A1, αSMA, VEGF-A, and VEGFR2 antibodies. In total, PEA material of 8 patients was evaluated and similar lesions were observed. Scale bar = 100 µm; purple arrow, CD31^+^ recanalizing vessel; red arrow, CD31^+^COL15A1^+^ recanalizing vessel; arrowhead, VEGFR2^+^ cells in media of recanalizing vessel, potentially smooth muscle cells. PA, pulmonary artery; PEA, pulmonary endarterectomy; MTC, Masson trichrome; αSMA, alpha smooth muscle actin; VEGF-A, vascular endothelial growth factor A; VEGFR2, vascular endothelial growth factor receptor 2.

**Figure 4:**
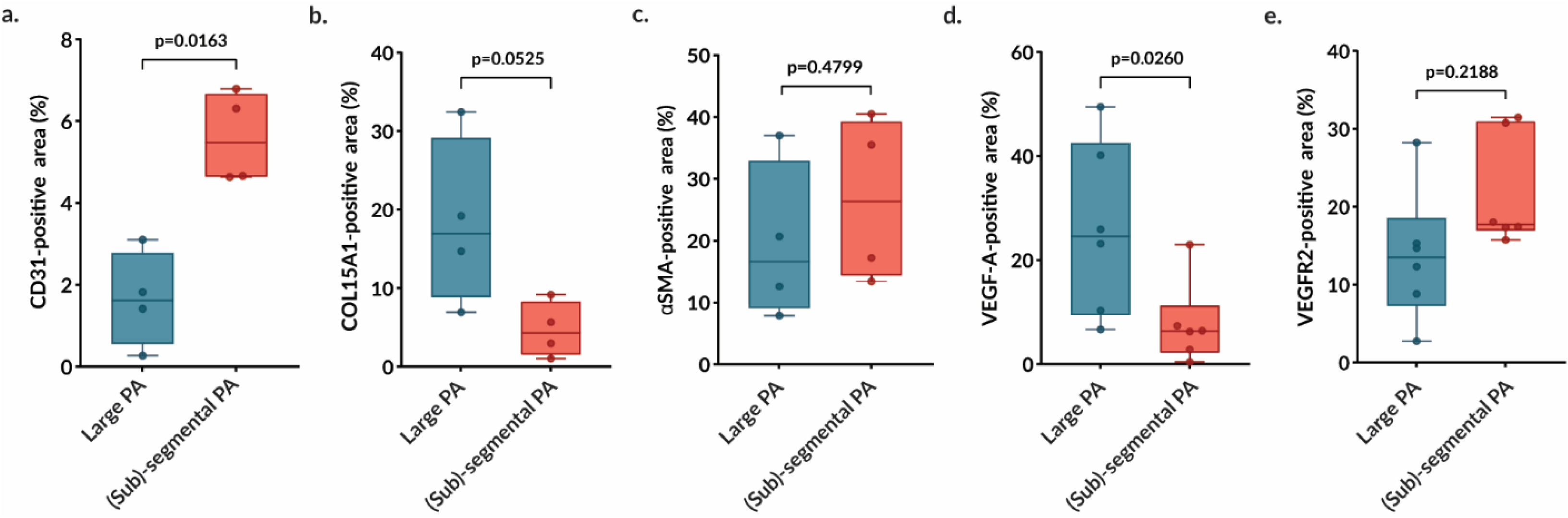
Expression profile of endothelial and angiogenic signaling markers in large and (sub)-segmental pulmonary arterial lesions of CTEPH patients. DAB-positive regions of CD31 (n=4), COL15A1 (n=4), VEGF-A (n=6), VEGFR2 (n=6) and αSMA (n=4) expression in large and (sub)-segmental pulmonary artery tissue were quantified using QuPath software. Due to technical reasons, patient cohort was reduced to n=4 for CD31,COL15A1 and αSMA, and n=6 for VEGF-A and VEGFR2. Results are expressed as median (25^th^-75^th^ interquartile range). Comparisons by paired Student and Wilcoxon t-test. PA, pulmonary artery; VEGF-A, vascular endothelial growth factor A; VEGFR2, vascular endothelial growth factor receptor 2; αSMA, alpha smooth muscle actin.

### In vitro angiogenic capacity of ECs from large and (sub)-segmental pulmonary arterial lesions

Given the differences observed in the expression of VEGF-A/VEGFR2 signaling pathway between large and (sub)-segmental pulmonary arterial lesions, we sought to determine whether the *in vitro* angiogenic properties of large and (sub)-segmental pulmonary arterial lesions would be differentially affected. Therefore, PAECs isolated from large and (sub)- segmental pulmonary arterial lesions were phenotyped, as illustrated in **Figure 5**. Both exhibit a typical “cobblestone” morphology and form a homogeneous monolayer (**Figure 5a & 5b**). Both large and (sub)-segmental PAECs express CD31 (**Figure 5c & 5d)** and VE-cadherin (**Figure 5e & 5f)** on their surface, vWF (**Figure 5g & 5h)** in their cytoplasm and take up acetylated LDL (**Figure 5i & 5j)** via endothelium-specific scavenger receptors. *In vitro* angiogenic capacity was assessed by performing migration and tube formation assays on PAECs from both large and (sub)-segmental pulmonary arterial lesions. Wound closure time (**Figure 6a)** and averaged migration area did not differ significantly over time (6 h: 1.29 ± 0.37 mm^2^/h vs. 1.32 ± 0.35 mm^2^/h; p = 0.99; 12 h: 2.25 ± 0.27 mm^2^/h vs. 2.35 ± 0.24 mm^2^/h; p = 0.93; 18 h: 2.45 ± 0.11 mm^2^/h vs. 2.56 ± 0.22 mm^2^/h; p = 0.75; and 20 h: 2.48 ± 0.12 mm^2^/h vs. 2.56 ± 0.22 mm^2^/h; p = 0.90) between large and (sub)-segmental PAECs (**Figure 6b**). Similarly, migration rate of large and (sub)-segmental PAECs was not significantly different (34 ± 3.4 mm^2^/h vs. 35 ± 3.7 mm^2^/h; p = 0.34) (**Figure 6c**). The average number of nodes (533 ± 234 vs. 484 ± 165; p = 0.49), segments 187 ± 95 vs. 165 ± 66; p = 0.45), junctions (152 ± 66 vs. 139 ± 46; p = 0.51) and branches (83 ± 10 vs. 83 ± 9; p = 0.90) did not differ significantly between large and (sub)-segmental PAECs (**Figure 7a-e**). These results suggest that *in vitro* angiogenic capacities are not different between ECs from large and (sub)-segmental pulmonary lesions of CTEPH patients.

**Figure 5:**
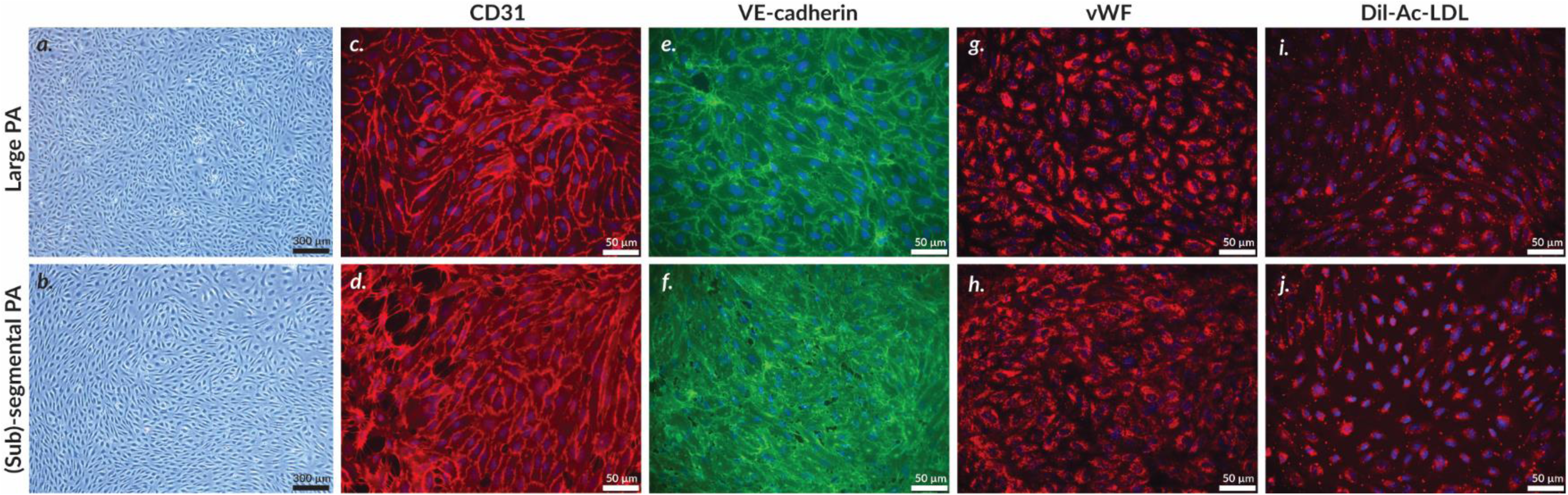
Phenotyping of PAECs from large and (sub)-segmental pulmonary arterial lesions of CTEPH patients. Primary culture of large and (sub)-segmental PAECs isolated from PEA material **(a,b)**. PAECs were characterized for endothelial phenotype by cell surface markers CD31**(c,d)** and VE-cadherin **(e,f)**, and intracellular endothelial marker vWF **(g,h)**. Endothelial phenotype was further confirmed by scavenger receptor capacity to internalize acetylated low-density-lipoprotein **(i,j)**. Nuclei were counterstained with DAPI (blue). Experiments were performed between passages 3 and 7. Scale = 300 µm (**a,b**) and 50 µm (**c-j**). PAECs, pulmonary artery endothelial cells; PA, pulmonary artery; PEA, pulmonary endarterectomy; vWF, von Willebrand factor.

**Figure 6:**
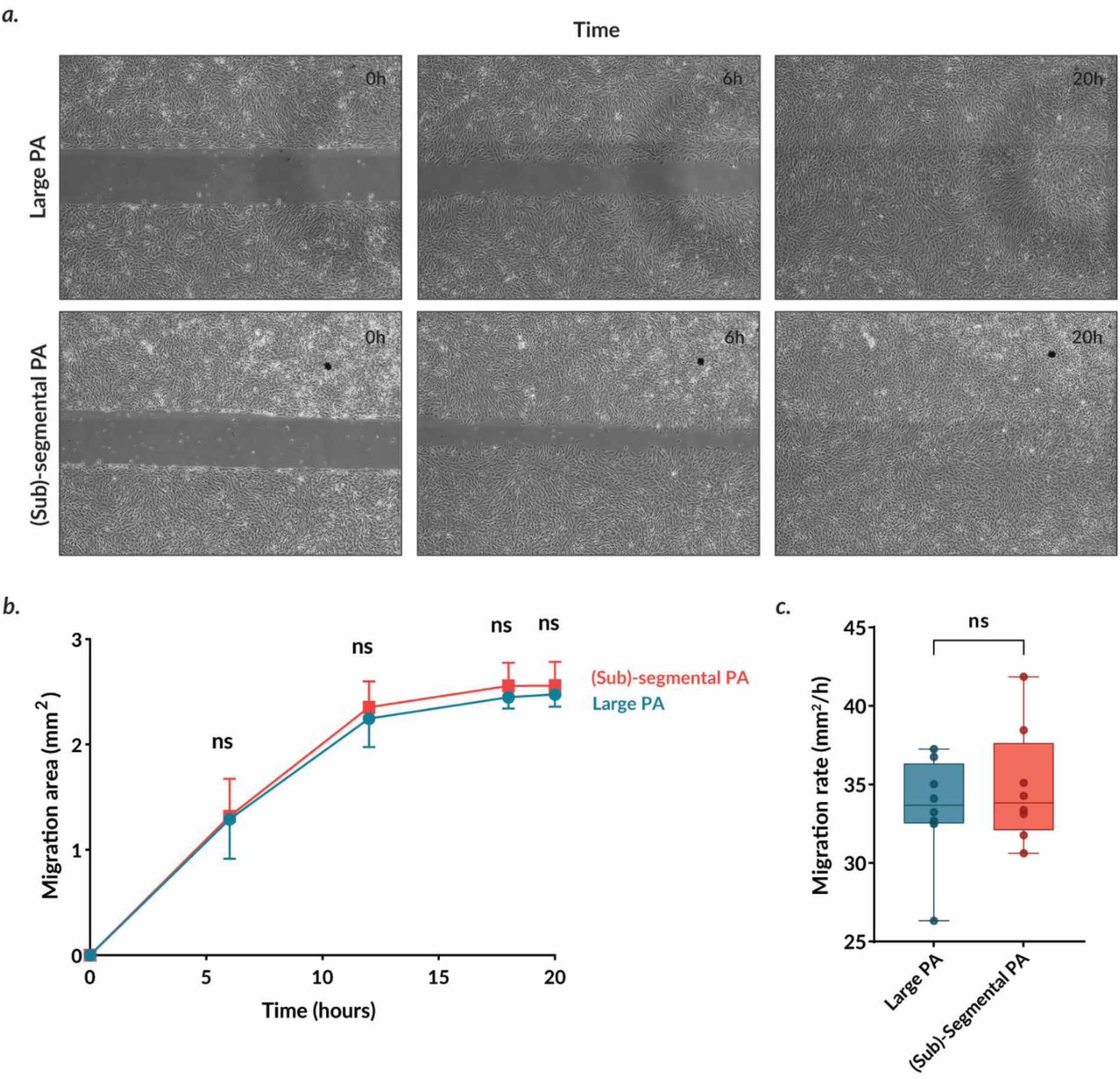
Migration capacity of PAECs isolated from large and (sub)-segmental pulmonary arterial lesions of CTEPH patients. Representative pictures of wound closure at baseline, 6 hours and 20 hours of large and (sub)-segmental PAECs **(a)**. Migration capacity of large and (sub)-segmental PAECs was assessed by measuring the gap area over time. Representative time curve of mean migration area (mm^2^) for large and (sub)-segmental PAECs **(b)**. Migration rate (mm^2^/h) of large and (sub)-segmental PAECs was assessed by measuring the area under the curve of the migration area **(c)**. Two independent experiments per patient (n=8) were carried out between passages 3 and 8 and expressed as median (25^th^-75^th^ interquartile range). Comparisons by two-way ANOVA and post-hoc Šídák’s test (**b**) and paired Student t-test (**c**). ns, non significant; PAECs, pulmonary artery endothelial cells; PA, pulmonary artery.

**Figure 7:**
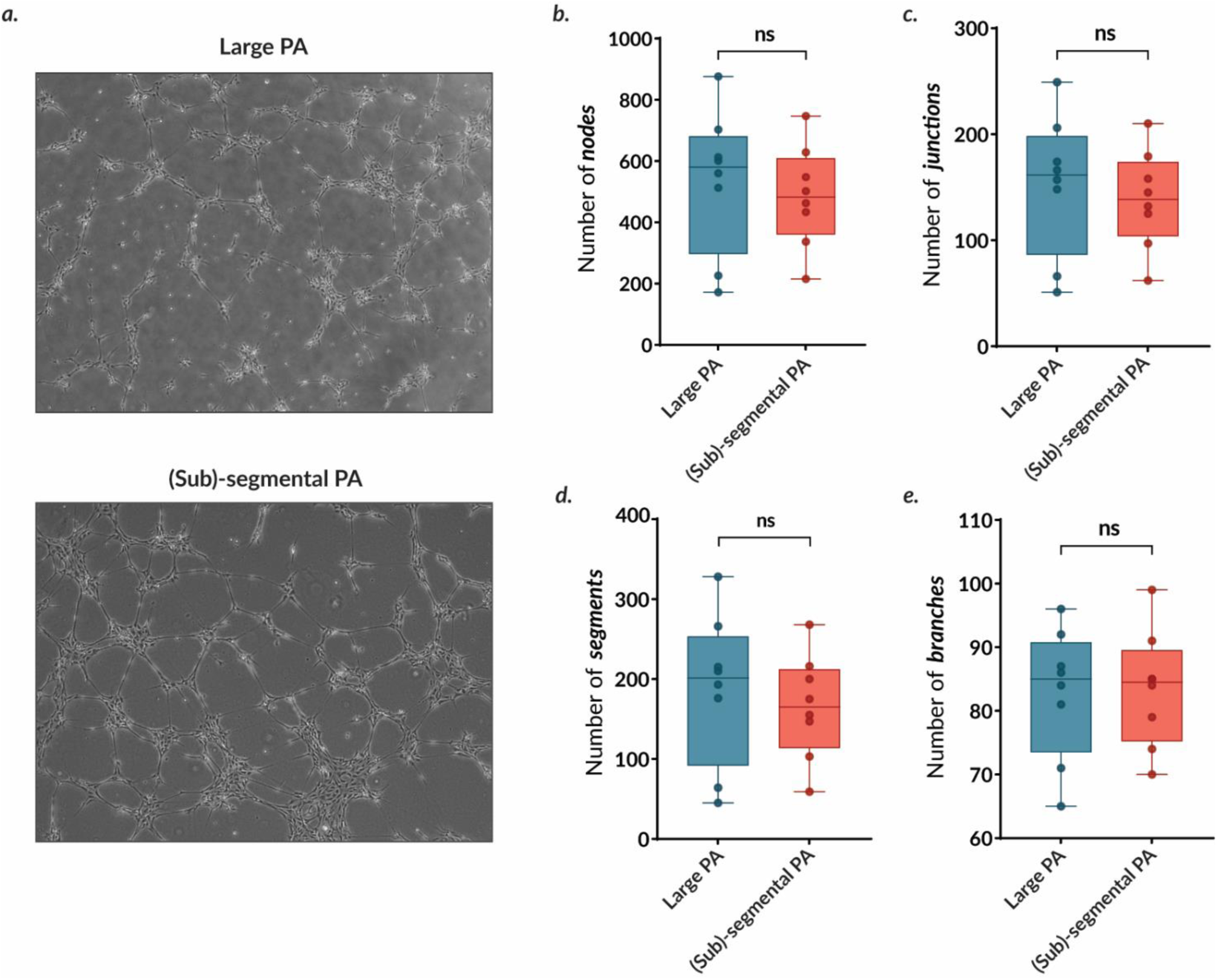
Tube formation capacity of PAECs isolated from large and (sub)-segmental pulmonary arterial lesions of CTEPH patients. Representative pictures of tube formation capacity of large and (sub)-segmental PAECs **(a)**. Tube formation capacity was assessed by counting the number of nodes **(b)**, junctions **(c)**, segments **(d)**, and branches **(e)**. Two independent experiments per patient (n=8) were carried out in triplicate between passages 3 and 7 and expressed as median (25^th^-75^th^ interquartile range). Comparisons by paired Student t-test (**b-e**). ns, non significant; PAECs, pulmonary artery endothelial cells; PA, pulmonary artery.

### Barrier function of ECs from large and (sub)-segmental pulmonary arteries

Knowing that various vascular pathologies are associated with impaired barrier function (45) and inflammatory cell infiltration has been observed in pulmonary arterial lesions from CTEPH patients (19), we sought to investigate whether *in vitro* endothelial barrier function might be differentially affected in large and (sub)-segmental pulmonary arterial lesions from CTEPH patients. BSA-FITC leakage through the endothelial monolayer over time did not differ significantly between PAECs from large and (sub)-segmental pulmonary lesions, (30 min: 0.26 ± 0.12 µg/mL vs. 0.25 ± 0.08 µg/mL; p > 0.99; 1 h: 0.46 ± 0.18 µg/mL vs. 0.49 ± 0.12 µg/mL; p > 0.99; 2 h: 0.81 ± 0.34 µg/mL vs. 0.84 ± 0.21 µg/mL; p > 0.99; 4 h: 1.18 ± 0.44 µg/mL vs. 1.32 ± 0.29 µg/mL; p = 0.96; **Figure 8a**). Similarly, BSA-leakage rate does not significantly differ between large and (sub)-segmental PAECs (1.53 µg/mL/h [1.30-1.75 µg/mL/h] vs. 1.60 µg/mL/h [1.48-2.04 µg/mL/h], median [interquartile range]; p = 0.74; **Figure 8b**). These results suggest that the endothelial barrier function is similarly affected in large and (sub)-segmental pulmonary lesions of CTEPH patients.

**Figure 8:**
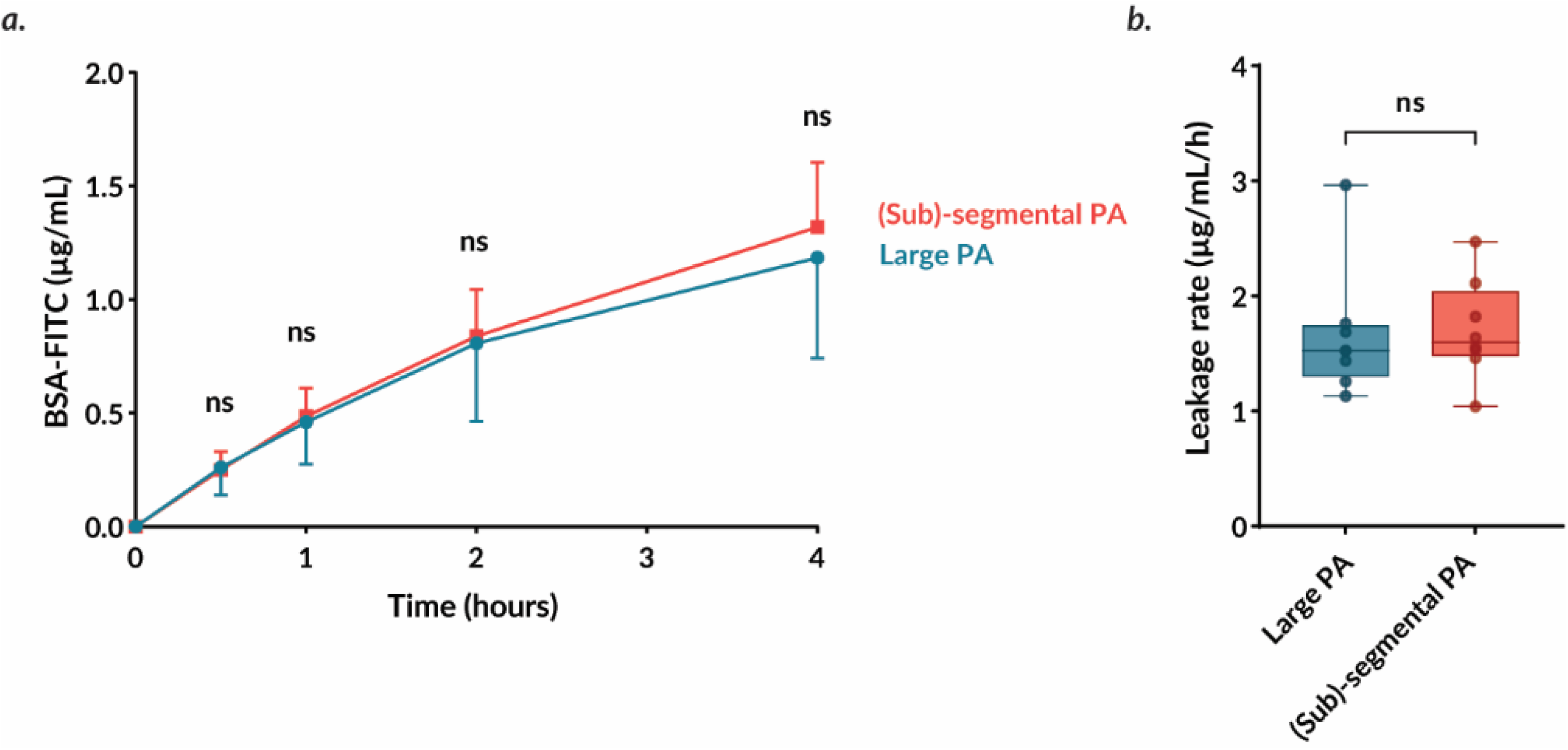
Barrier function of PAECs isolated from large and (sub)-segmental pulmonary arterial lesions of CTEPH patients. Leakage of BSA-FITC (µg/mL) through large and (sub)-segmental PAECs monolayer was assessed over time **(a)**. Averaged BSA-FITC leakage rate (µg/mL/h) through large and (sub)-segmental PAECs monolayer was evaluated by measuring the area under the curve **(b)**. Two independent experiments per patient (n=8) were carried out in duplicate between passages 3 and 6 and expressed as median (25^th^-75^th^ interquartile range). Comparisons by two-way ANOVA and post-hoc Šídák’s test **(a)** and paired Wilcoxon t-test **(b)**. ns, non significant; PAECs, pulmonary artery endothelial cells; PA, pulmonary artery.

## DISCUSSION

In this study, we observed differences in *in situ* angiogenic cues of the VEGF-A/VEGFR2 axis, whereas *in vitro* angiogenic capacities and endothelial barrier function did not differ significantly according to the location (large *vs.* (sub)-segmental) along the pulmonary vascular lesion tree. It is noteworthy that only a subset of vessels recanalizing (sub)-segmental pulmonary arterial lesions exhibit expression of COL15A1, a marker of systemic endothelial cells (38). This observation suggests that the source of endothelial cells responsible for the recanalization of pulmonary arterial lesions in CTEPH patients is likely to be heterogeneous, potentially involving ECs from both the systemic and pulmonary circulation or even progenitor ECs.

The present study revealed the presence of muscle and collagen fibers in both large and (sub)- segmental pulmonary arterial lesions. In large pulmonary arterial lesions, numerous myofibroblast-like cells are present in the neointima, which provides evidence of intense vascular remodeling, as previously described (17,18,37). The presence of αSMA-positive myofibroblast-like cells may result from endothelial-to-mesenchymal transition (endoMT), which could further contribute to the formation of fibro-thrombotic material obstructing pulmonary arteries in CTEPH (46,47). Interestingly, Bocheneck *et al.* have previously shown that TGFβ1 signaling, known to promote endoMT, impedes venous thrombus resolution potentially contributing to the pathogenesis of CTEPH (48,49).

COL15A1 is a member of the multiplexin superfamily of collagens, produced by fibroblasts, smooth muscle cells and endothelial cells (50,51). Its primary function is to organize, stabilize and integrate the basement membrane with underlying connective tissue (50). Recently, Schupp et al. identified COL15A1 as a specific marker of systemic endothelial cells in human lungs using single-cell RNA sequencing (38). In (sub)-segmental recanalyzed pulmonary arterial lesions, we observed the presence of both CD31-positive/COL15A1-positive cells, and CD31-positive/COL15A1-negative cells. Additionally, we found that PAECs lining the pulmonary arterial lumen of large vessels exhibit a CD31-positive/COL15A1-negative pattern. These findings suggest that adjacent endothelial cells are replicating to reform a neointima, likely in conjunction with progenitor cells together with progenitor cells (26,52,53), in partially occluded regions of large pulmonary arteries. Conversely, neovessels recanalizing fibro-thrombotic lesions occluding (sub)-segmental pulmonary arteries may originate from systemic bronchopulmonary anastomoses (54). Surprisingly, within large pulmonary arterial lesions, CD31-negative myofibroblast-like cells do express COL15A1. This suggests either involvement of endoMT as abovementioned, or smooth muscle dedifferentiation, which is often observed in neointima formation.

Given the pivotal role of angiogenesis in thrombus resolution (36) and in CTEPH progression, as recently demonstrated in our CTEPH rabbit model (35), as well as its potential impact on outcome of CTEPH patients (19), it is not surprising that the angiogenic VEGF-A/VEGFR2 axis emerged as a potential contributor.

VEGF-A is expressed in both large and (sub)-segmental pulmonary arterial lesions, with a greater level observed in large pulmonary arterial lesions. This may appear counterintuitive when considering the large presence of recanalizing vessels in (sub)-segmental pulmonary arterial lesions. Within (sub)-segmental pulmonary arterial lesions, CD31-positive cells lining recanalizing vessels do not express VEGF-A, suggesting that alternative ligands may be involved. These include VEGF-C or VEGF-D, although their binding affinity is weaker compared to VEGF-A, or VEGF-B and placenta growth factor (PIGF) via VEGFR1 (55). In contrast, Bochenek et al. previously observed that the endothelium of vessels recanalizing lesions do express VEGF (37), although VEGF type was not identified. Finally, VEGFR2 is expressed by myofibroblast-like and medial smooth muscle cells in pulmonary arterial lesions, with no notable distinction between large and (sub)-segmental pulmonary arterial lesions. Accordingly, Bochenek et al. also observed VEGFR2 expression in spindle-shaped cells within organized thrombi and myofibroblasts (37).

To further elucidate the differential features of the angiogenic VEGF-A/VEGFR2 axis observed between large and (sub)-segmental pulmonary arterial lesions, we aim to investigate their *in vitro* angiogenic properties, knowing that migration and tube formation capacity of PAECs from PEA material can be impaired, in comparison with control PAECs from healthy donors (35). However, no discernable distinction was observed between PAECs from large and (sub)- segmental lesions. These findings are not consistent with the CD31-positive high content and the numerous recanalizing vessels observed in (sub)-segmental pulmonary arterial lesions. It should be noted, however, that the PAECs were isolated from PEA material of CTEPH patients at relatively advanced disease stages. Therefore, it may be beneficial to consider alternative approaches, such as gaining cells at diagnosis via right heart catheterization (40) to overcome this limitation. Furthermore, angiogenic capacities are assessed using Matrigel^©^, which has a markedly different composition from the *in vivo* extracellular matrix.

Furthermore, an imbalance of the VEGF-A/VEGFR2 axis can also affect vascular permeability and endothelial cell function (55). In vascular pathology such as CTEPH, the endothelium plays a pivotal role in governing the movement of circulating proteins and cells between the bloodstream and the underlying tissues. As previously demonstrated, inflammatory stimuli can compromise endothelial integrity in the context of pulmonary hypertension (44). Additionally, the accumulation of inflammatory cells, including T-lymphocytes, macrophage and neutrophils, has been observed in PEA material of CTEPH patients (19). Moreover, ischemia and alterations in shear stress have been demonstrated to affect endothelial barrier function (57–59). However, no significant difference in barrier function was observed between large and (sub)-segmental PAECs, suggesting that endothelial barrier function is similarly impaired in both vascular beds.

It should be noted that the study is not without limitations. The authors are aware that the *in vitro* results are based on primary cells maintained in culture. Despite the use of a standardized isolation protocol, it cannot be excluded that a biased selection of the most robust cell clusters may have occurred. As previously stated, cells were isolated from PEA material obtained from patients who had already reached advanced stages of the disease. This does not illustrate the potential of the cells at early phases, following an acute episode of pulmonary embolism. Finally, the utilization of synthetic hydrogel in simplified two-dimensional *in vitro* assays, in the absence of blood flow and shear stress conditions, does not fully replicate the *in vivo* conditions and circumvents interactions between cells and the extracellular matrix.

In conclusion, both systemic and pulmonary endothelial cells contribute to the recanalization of fibro-thrombotic pulmonary vascular lesions in CTEPH patients. Despite the existence of different *in situ* angiogenic cues in the VEGF-A/VEGFR2 axis according to the location along the pulmonary vascular tree, *in vitro* angiogenic capacities and barrier function remain unchanged. This suggests that simplified *in vitro* assays are probably not completely adequate to explain all *in situ* features. Consequently, the emergence of more sophisticated *in vitro* systems such as organoids and microfluidic devices, in which cells and organ-derived ECM could be combined, should contribute to elucidating the mechanisms potentially involved.

## Ethical approval

The study protocol was approved by the Institutional Ethics Committee of the University Hospital of Leuven (S57114).

## Guarantor

not applicable

## Author contributions

RQ conceived and designed the study, contributed to data interpretation, reviewed the manuscript; JV and LW collected the data, performed data analysis and interpretation; drafted the manuscript. AW, RS, ND and LA collected data and reviewed the manuscript. TV and MD collected patient material and reviewed the manuscript; all authors approved the final version.

## REFERENCES

1. Delcroix M, Torbicki A, Gopalan D, Sitbon O, Klok FA, Lang I, et al. ERS statement on chronic thromboembolic pulmonary hypertension. Eur Respir J. 2021;57:2002828.

2. Humbert M, Kovacs G, Hoeper MM, Badagliacca R, Berger RMF, Brida M, et al. 2022 ESC/ERS Guidelines for the diagnosis and treatment of pulmonary hypertension. Eur Respir J. 2022;63:2200879.

3. Humbert M, Kovacs G, Hoeper MM, Badagliacca R, Berger RMF, Brida M, et al. 2022 ESC/ERS Guidelines for the diagnosis and treatment of pulmonary hypertension. Eur Heart J. 2022;43:3618–731.

4. Klok FA, Couturaud F, Delcroix M, Humbert M. Diagnosis of chronic thromboembolic pulmonary hypertension after acute pulmonary embolism. Eur Respir J. 2020;55:2000189.

5. Simonneau G, Torbicki A, Dorfmüller P, Kim N. The pathophysiology of chronic thromboembolic pulmonary hypertension. Eur Respir Rev. 2017;26:160112.

6. Kim NH, Delcroix M, Jais X, Madani MM, Matsubara H, Mayer E, et al. Chronic thromboembolic pulmonary hypertension. Eur Respir J. 2019;53:1801915.

7. Galie N. Pulmonary Microvascular Disease in Chronic Thromboembolic Pulmonary Hypertension. Proc Am Thorac Soc. 2006;3:571–6.

8. Moser KM, Bloor CM. Pulmonary vascular lesions occurring in patients with chronic major vessel thromboembolic pulmonary hypertension. Chest. 1993;103:685–92.

9. Dorfmüller P, Günther S, Ghigna MR, Thomas de Montpréville V, Boulate D, Paul JF, et al. Microvascular disease in chronic thromboembolic pulmonary hypertension: a role for pulmonary veins and systemic vasculature. Eur Respir J. 2014;44:1275–88.

10. Jenkins D. Pulmonary endarterectomy: the potentially curative treatment for patients with chronic thromboembolic pulmonary hypertension. Eur Respir Rev. 2015;24:263–71.

11. Elwing JM, Vaidya A, Auger WR. Chronic Thromboembolic Pulmonary Hypertension: An Update. Clin Chest Med. 2018;39:605–20.

12. Kim NH, Lang IM. Risk factors for chronic thromboembolic pulmonary hypertension. Eur Respir Rev. 2012;21:27–31.

13. Lang IM, Pesavento R, Bonderman D, Yuan JXJ. Risk factors and basic mechanisms of chronic thromboembolic pulmonary hypertension: a current understanding. Eur Respir J. 2013;41:462–8.

14. Manz XD, Szulcek R, Pan X, Symersky P, Dickhoff C, Majolée J, et al. Epigenetic Modification of the von Willebrand Factor Promoter Drives Platelet Aggregation on the Pulmonary Endothelium in Chronic Thromboembolic Pulmonary Hypertension. Am J Respir Crit Care Med. 2022;205:806– 18.

15. Manz XD, Bogaard HJ, Aman J. Regulation of VWF (Von Willebrand Factor) in Inflammatory Thrombosis. Arterioscler Thromb Vasc Biol. 2022;42:1307–20.

16. Arbustini E, Morbini P, D’Armini AM, Repetto A, Minzioni G, Piovella F, et al. Plaque composition in plexogenic and thromboembolic pulmonary hypertension: the critical role of thrombotic material in pultaceous core formation. Heart Br Card Soc. 2002;88:177–82.

17. Bernard J, Yi ES. Pulmonary thromboendarterectomy: a clinicopathologic study of 200 consecutive pulmonary thromboendarterectomy cases in one institution. Hum Pathol. 2007;38:871–7.

18. Blauwet LA, Edwards WD, Tazelaar HD, McGregor CGA. Surgical pathology of pulmonary thromboendarterectomy: a study of 54 cases from 1990 to 2001. Hum Pathol. 2003;34:1290–8.

19. Quarck R, Wynants M, Verbeken E, Meyns B, Delcroix M. Contribution of inflammation and impaired angiogenesis to the pathobiology of chronic thromboembolic pulmonary hypertension. Eur Respir J. 2015;46:431–43.

20. Kimura H, Okada O, Tanabe N, Tanaka Y, Terai M, Takiguchi Y, et al. Plasma Monocyte Chemoattractant Protein-1 and Pulmonary Vascular Resistance in Chronic Thromboembolic Pulmonary Hypertension. Am J Respir Crit Care Med. 2001;164:319–24.

21. Quarck R, Nawrot T, Meyns B, Delcroix M. C-Reactive Protein. J Am Coll Cardiol. 2009;53:1211– 8.

22. Langer F, Schramm R, Bauer M, Tscholl D, Kunihara T, Schäfers HJ. Cytokine Response to Pulmonary Thromboendarterectomy. CHEST. 2004;126:135–41.

23. Wynants M, Vengethasamy L, Ronisz A, Meyns B, Delcroix M, Quarck R. NF-κB pathway is involved in CRP-induced effects on pulmonary arterial endothelial cells in chronic thromboembolic pulmonary hypertension. Am J Physiol-Lung Cell Mol Physiol. 2013;305:L934–42.

24. Wynants M, Quarck R, Ronisz A, Alfaro-Moreno E, Raemdonck DV, Meyns B, et al. Effects of C-reactive protein on human pulmonary vascular cells in chronic thromboembolic pulmonary hypertension. Eur Respir J. 2012;40:886–94.

25. Firth AL, Yao W, Ogawa A, Madani MM, Lin GY, Yuan JXJ. Multipotent mesenchymal progenitor cells are present in endarterectomized tissues from patients with chronic thromboembolic pulmonary hypertension. Am J Physiol Cell Physiol. 2010;298:C1217–1225.

26. Yao W, Firth AL, Sacks RS, Ogawa A, Auger WR, Fedullo PF, et al. Identification of putative endothelial progenitor cells (CD34+CD133+Flk-1+) in endarterectomized tissue of patients with chronic thromboembolic pulmonary hypertension. Am J Physiol Lung Cell Mol Physiol. 2009;296:L870–878.

27. Waltham M, Burnand KG, Collins M, McGuinness CL, Singh I, Smith A. Vascular endothelial growth factor enhances venous thrombus recanalisation and organisation. Thromb Haemost. 2003;89:169–76.

28. Waltham M, Burnand K, Fenske C, Modarai B, Humphries J, Smith A. Vascular endothelial growth factor naked DNA gene transfer enhances thrombus recanalization and resolution. J Vasc Surg. 2005;42:1183–9.

29. Modarai B, Humphries J, Gossage JA, Waltham M, Burnand KG, Kanaganayagam GS, et al. Adenovirus-Mediated VEGF Gene Therapy Enhances Venous Thrombus Recanalization and Resolution. Arterioscler Thromb Vasc Biol. 2008;28:1753–9.

30. Alias S, Redwan B, Panzenböck A, Winter MP, Schubert U, Voswinckel R, et al. Defective Angiogenesis Delays Thrombus Resolution: A Potential Pathogenetic Mechanism Underlying Chronic Thromboembolic Pulmonary Hypertension. Arterioscler Thromb Vasc Biol. 2014;34:810– 9.

31. Evans CE, Grover SP, Humphries J, Saha P, Patel AP, Patel AS, et al. Antiangiogenic Therapy Inhibits Venous Thrombus Resolution. Arterioscler Thromb Vasc Biol. 2014 Mar;34(3):565–70.

32. Hobohm L, Kölmel S, Niemann C, Kümpers P, Krieg VJ, Bochenek ML, et al. Role of angiopoietin- 2 in venous thrombus resolution and chronic thromboembolic disease. Eur Respir J. 2021;58:2004196.

33. Hosokawa K, Ishibashi-Ueda H, Kishi T, Nakanishi N, Kyotani S, Ogino H. Histopathological Multiple Recanalized Lesion Is Critical Element of Outcome After Pulmonary Thromboendarterectomy. Int Heart J. 2011;52:377–81.

34. Viswanathan G, Kirshner HF, Nazo N, Ali S, Ganapathi A, Cumming I, et al. Single-Cell Analysis Reveals Distinct Immune and Smooth Muscle Cell Populations that Contribute to Chronic Thromboembolic Pulmonary Hypertension. Am J Respir Crit Care Med. 2023;207:1358–75.

35. Quarck R, Tielemans B, Willems L, Stoian L, Ronisz A, Wagenaar A, et al. Impairment of angiogenesis-driven clot resolution is a key event in the progression to CTEPH: validation in a novel rabbit model. Arterioscler Thromb Vasc Biol. 2023;43):1308–21.

36. Willems L, Kurakula K, Verhaegen J, Klok FA, Delcroix M, Goumans MJ, et al. Angiogenesis in Chronic Thromboembolic Pulmonary Hypertension: A Janus-Faced Player? Arterioscler Thromb Vasc Biol. 2024;44:794–806 2.

37. Bochenek ML, Rosinus NS, Lankeit M, Hobohm L, Bremmer F, Schütz E, et al. From thrombosis to fibrosis in chronic thromboembolic pulmonary hypertension. Thromb Haemost. 2017;117:769– 83.

38. Schupp JC, Adams TS, Cosme C, Raredon MSB, Yuan Y, Omote N, et al. Integrated Single-Cell Atlas of Endothelial Cells of the Human Lung. Circulation. 2021;144:286–302.

39. Quarck R, Wynants M, Ronisz A, Sepulveda MR, Wuytack F, Van Raemdonck D, et al. Characterization of proximal pulmonary arterial cells from chronic thromboembolic pulmonary hypertension patients. Respir Res. 2012;13:27.

40. Tielemans B, Stoian L, Wagenaar A, Leys M, Belge C, Delcroix M, et al. Incremental Experience in In Vitro Primary Culture of Human Pulmonary Arterial Endothelial Cells Harvested from Swan-Ganz Pulmonary Arterial Catheters. Cells. 2021;10:3229.

41. Vengethasamy L, Hautefort A, Tielemans B, Belge C, Perros F, Verleden S, et al. BMPRII influences the response of pulmonary microvascular endothelial cells to inflammatory mediators. Pflüg Arch - Eur J Physiol. 2016;468:1969–83.

42. Tielemans B, Wagenaar A, Belge C, Delcroix M, Quarck R. Pulmonary arterial hypertension drugs can partially restore altered angiogenic capacities in bmpr2-silenced human lung microvascular endothelial cells. Pulm Circ. 2023;13:e12293.

43. 43. Carpentier G. Angiogenic Analyzer [Internet]. Available from: http://image.bio.methods.free.fr/ImageJ/?Angiogenesis-Analyzer-for-ImageJ.

44. Tielemans B, Stoian L, Gijsbers R, Michiels A, Wagenaar A, Farre Marti R, et al. Cytokines trigger disruption of endothelium barrier function and p38 MAP kinase activation in BMPR2-silenced human lung microvascular endothelial cells. Pulm Circ. 2019;9:2045894019883607.

45. Park-Windhol C, D’Amore PA. Disorders of Vascular Permeability. Annu Rev Pathol. 2016;11:251–81.

46. Sakao S, Hao H, Tanabe N, Kasahara Y, Kurosu K, Tatsumi K. Endothelial-like cells in chronic thromboembolic pulmonary hypertension: crosstalk with myofibroblast-like cells. Respir Res. 2011;12:109.

47. Arciniegas E, Frid MG, Douglas IS, Stenmark KR. Perspectives on endothelial-to-mesenchymal transition: potential contribution to vascular remodeling in chronic pulmonary hypertension. Am J Physiol-Lung Cell Mol Physiol. 2007;293:L1–8.

48. Bochenek ML, Leidinger C, Rosinus NS, Gogiraju R, Guth S, Hobohm L, et al. Activated Endothelial TGFβ1 Signaling Promotes Venous Thrombus Nonresolution in Mice Via Endothelin-1. Circ Res. 2020;126:162–81.

49. Bochenek ML, Saar K, Nazari-Jahantigh M, Gogiraju R, Wiedenroth CB, Münzel T, et al. Endothelial Overexpression of TGF-β-Induced Protein Impairs Venous Thrombus Resolution. JACC Basic Transl Sci. 2024;9:100–16.

50. Myers JC, Dion AS, Abraham V, Amenta PS. Type XV collagen exhibits a widespread distribution in human tissues but a distinct localization in basement membrane zones. Cell Tissue Res. 1996;286:493–505.

51. Prockop DJ, Kivirikko KI. COLLAGENS: Molecular Biology, Diseases, and Potentials for Therapy. Annu Rev Biochem. 1995;64:403–34.

52. Nicklas JM, Gordon AE, Henke PK. Resolution of Deep Venous Thrombosis: Proposed Immune Paradigms. Int J Mol Sci. 2020;21:2080.

53. Alvarez DF, Huang L, King JA, ElZarrad MK, Yoder MC, Stevens T. Lung microvascular endothelium is enriched with progenitor cells that exhibit vasculogenic capacity. Am J Physiol-Lung Cell Mol Physiol. 2008;294:L419–30.

54. Dorfmüller P, Günther S, Ghigna MR, Montpréville VT de, Boulate D, Paul JF, et al. Microvascular disease in chronic thromboembolic pulmonary hypertension: a role for pulmonary veins and systemic vasculature. Eur Respir J. 2014;44:1275–88.

55. Shibuya M, Claesson-Welsh L. Signal transduction by VEGF receptors in regulation of angiogenesis and lymphangiogenesis. Exp Cell Res. 2006;312:549–60.

56. Tura-Ceide O, Smolders VFED, Aventin N, Morén C, Guitart-Mampel M, Blanco I, et al. Derivation and characterisation of endothelial cells from patients with chronic thromboembolic pulmonary hypertension. Sci Rep. 2021;11:18797.

57. Nagy JA, Benjamin L, Zeng H, Dvorak AM, Dvorak HF. Vascular permeability, vascular hyperpermeability and angiogenesis. Angiogenesis. 2008;11(2):109–19.

58. Weis SM. Vascular permeability in cardiovascular disease and cancer. Curr Opin Hematol. 2008;15:243–9.

59. Paszkowiak JJ, Dardik A. Arterial wall shear stress: observations from the bench to the bedside. Vasc Endovascular Surg. 2003;37:47–57.

