## Supplemental figures for "Endothelial features along the pulmonary vascular tree in chronic thromboembolic pulmonary hypertension: distinctive or shared facets?"

**SUPPLEMENTAL DATA**

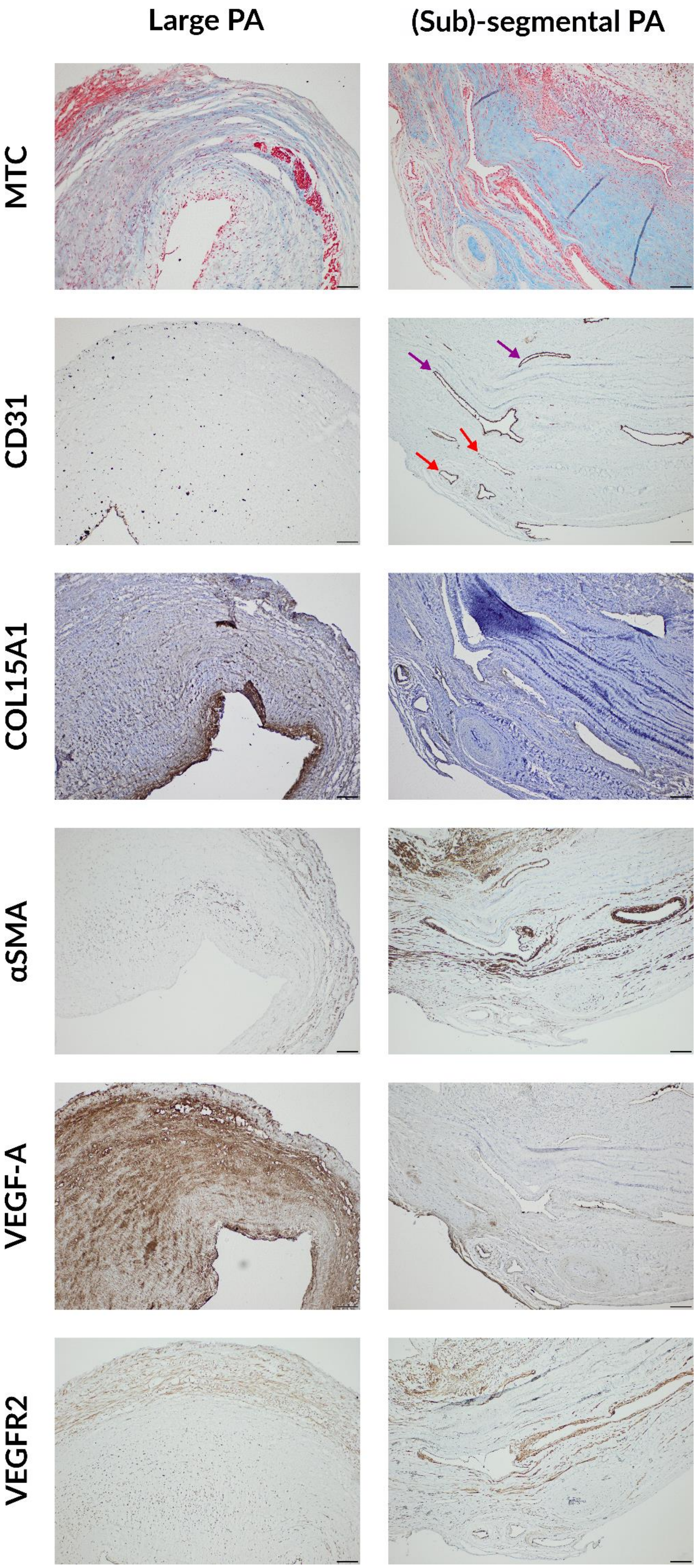
Large PA
(Sub)-segmental PA
MTC
CD31
COL15A1
αSMA
VEGF-A
VEGFR2

**Figure S1: Histological evaluation of large and (sub)-segmental pulmonary arterial lesions of CTEPH patients (1).** Large and (sub)-segmental pulmonary arterial lesions were stained with Masson's trichrome stain (collagen, blue; muscle fibers, red) and immunolabelled using COL15A1, CD31, αSMA, VEGF-A, and VEGFR2 antibodies. In total, PEA material of 8 patients was evaluated and similar lesions were observed. Scale bar = 100 µm*;* purple arrow, $CD31^+$ recanalizing vessels; red arrow, $CD31^+COL15A1^+$ recanalizing vessels. PA, pulmonary artery; MTC, Masson trichrome; αSMA, alpha smooth muscle actin; VEGF-A, vascular endothelial growth factor A; VEGFR2, vascular endothelial growth factor receptor 2.

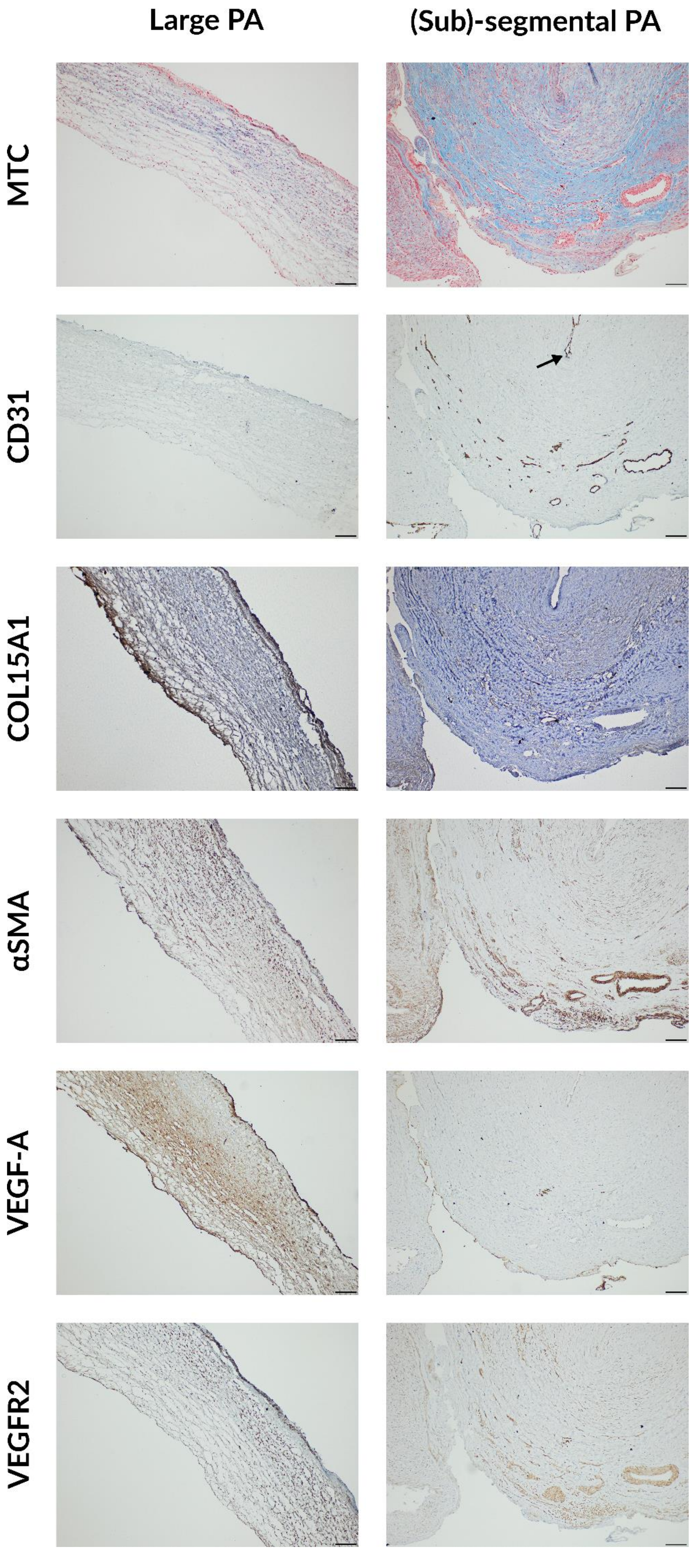

Large PA
(Sub)-segmental PA
MTC
CD31
COL15A1
αSMA
VEGF-A
VEGFR2

**Figure S2: Histological evaluation of large and (sub)-segmental pulmonary arterial lesions of CTEPH patients (2).** Large and (sub)-segmental pulmonary arterial lesions were stained with Masson's trichrome stain (collagen, blue; muscle fibers, red) and immunolabelled using COL15A1, CD31, αSMA, VEGF-A, and VEGFR2 antibodies. In total, PEA material of 8 patients was evaluated and similar lesions were observed. Scale bar = 100 µm; arrow, $CD31^{+}COL15A1^{-}$ cells lining the lumen of the obstructed segmental pulmonary arterial lesions. PA, pulmonary artery; MTC, Masson trichrome; αSMA, alpha smooth muscle actin; VEGF-A, vascular endothelial growth factor A; VEGFR2, vascular endothelial growth factor receptor 2.
